# Human Surfactant Protein A Alleviates SARS-CoV-2 Infectivity in Human Lung Epithelial Cells

**DOI:** 10.1101/2023.04.03.535215

**Authors:** Ikechukwu B Jacob, Amanda Gemmiti, Weichuan Xiong, Erin Reynolds, Brian Nicholas, Saravanan Thangamani, Hongpeng Jia, Guirong Wang

## Abstract

SARS coronavirus 2 (SARS-CoV-2) infects human angiotensin-converting enzyme 2 (hACE2)-expressing lung epithelial cells through its spike (S) protein. The S protein is highly glycosylated and could be a target for lectins. Surfactant protein A (SP-A) is a collagen-containing C-type lectin, expressed by mucosal epithelial cells and mediates its antiviral activities by binding to viral glycoproteins. This study examined the mechanistic role of human SP-A in SARS-CoV-2 infectivity. The interactions between human SP-A and SARS-CoV-2 S protein and hACE2 receptor, and SP-A level in COVID-19 patients were assessed by ELISA. The effect of SP-A on SARS-CoV-2 infectivity was analyzed by infecting human lung epithelial cells (A549-ACE2) with pseudoviral particles and infectious SARS-CoV-2 (Delta variant) pre-incubated with SP-A. Virus binding, entry, and infectivity were assessed by RT-qPCR, immunoblotting, and plaque assay. The results showed that human SP-A can bind SARS-CoV-2 S protein/RBD and hACE2 in a dose-dependent manner (p<0.01). Human SP-A inhibited virus binding and entry, and reduce viral load in lung epithelial cells, evidenced by the dose-dependent decrease in viral RNA, nucleocapsid protein, and titer (p<0.01). Increased SP-A level was observed in the saliva of COVID-19 patients compared to healthy controls (p<0.05), but severe COVID-19 patients had relatively lower SP-A levels than moderate COVID-19 patients (p<0.05). Therefore, SP-A plays an important role in mucosal innate immunity against SARS-CoV-2 infectivity by directly binding to the S protein and inhibiting its infectivity in host cells. SP-A level in the saliva of COVID-19 patients might serve as a biomarker for COVID-19 severity.

## Introduction

Nearly 7 million people have died due to coronavirus disease 2019 (COVID-19), with the United States reporting more deaths than any other country. COVID-19 is an infectious disease caused by severe acute respiratory syndrome coronavirus 2 (SARS-CoV-2)(1). Previous studies have shown that high morbidity and mortality following SARS-CoV-2 infection are predominantly due to a robust influx of inflammatory cells and cytokines into the lungs resulting in acute lung injury (ALI) and acute respiratory distress syndrome (ARDS)(2). SARS-CoV-2, like other coronaviruses, is an enveloped virus with several structural and non-structural proteins that facilitate its infectivity and pathogenicity in humans (3). Most important among the structural proteins for viral infectivity is the spike protein (S protein) which associates as a trimer on the viral envelope and is the basic unit through which the virus attaches to the host cellular receptor, human angiotensin-converting enzyme receptor 2 (hACE2), predominantly expressed on epithelial cells, including alveolar type II cells (ATII) in the lungs and in several other tissues (4). Each monomer of the S protein is composed of the S1 and S2 subunits. The S1 subunit contains the receptor-binding domain (RBD) which primarily binds to hACE2 while the S2 domain mediates the fusion of the viral and host cell membrane upon cleavage of the S protein subunit by the host transmembrane protease serine 2 (TMPRSS2) (5). Interestingly, as observed in most viral glycoproteins, the SARS-CoV-2 S protein is decorated with several N- and O-linked carbohydrate structures that have been demonstrated to protect it from antibody recognition (5, 6). While the presence of sugars on viral S protein can enable immune evasion, it may also enhance recognition by host innate immune carbohydrate-binding proteins (lectins), such as the human surfactant protein A (SP-A).

Human SP-A is a hydrophilic protein and belongs to the C-type lectin family of proteins (collectins) that surveys mucosal epithelial surfaces of the lungs, regions of the upper airway including laryngeal tissues, salivary glands, oral gingiva, and nasal mucosa, and bind to pathogen-associated molecular patterns (PAMPs) of most invading microbes (4, 7). Like other collectins, SP-A is composed of four functional domains among which is a carbohydrate recognition domain (CRD) that mediates Ca^2+^-dependent binding to sugars on microbial glycoproteins (7). As a pattern recognition molecule (PRM), SP-A, alongside a related human lung-associated collectin (SP-D), has been demonstrated to bind sugar moieties on viral surfaces and inhibit their infectivity (7). Furthermore, SP-A enhances viral aggregation, opsonization, and lysis while modulating inflammation by interacting with various types of receptors on innate immune cells (7, 8). Several studies have described the antiviral and immunomodulatory activities of SP-A in the context of respiratory syncytial virus (RSV), Influenza A virus (IAV), human coronavirus 229E (HCoV-229E), and HIV (9–12). Interestingly, a recent *in silico* analysis showed that SP-A could ligate the S protein with an affinity similar to the ACE2-spike interaction (13), suggesting that SP-A may have an implication in the pathogenesis of SARS-CoV-2 infection. Given that the effectiveness of the newly developed vaccines and therapeutics for COVID-19 is continuously being threatened by the frequent emergence of SARS-CoV-2 variants with unique changes in the spike epitope that facilitate immune escape, there is a considerable ongoing global effort to develop and improve antivirals and immunomodulatory agents (14–16).

In this study, we explored whether SP-A can bind SARS-CoV-2 S protein, RBD and hACE2, and inhibit viral entry in susceptible host cells. Our results revealed important information about the inhibitory role of human SP-A on SARS-CoV-2 infectivity and its mucosal innate immune response following SARS-CoV-2 infection. Moreover, the level of SP-A in the saliva of COVID-19 patients was also assessed. These findings contribute to our understanding of the role of human SP-A in SARS-CoV-2 induced pathogenesis and highlight SP-A as an important host protein that could serve as a biomarker for COVID-19 severity as well as a potential therapeutic component.

## Materials and Methods

### Human SP-A Protein

Native human SP-A (hSP-A) was isolated and purified from bronchoalveolar lavage fluid (BALF) of alveolar proteinosis patients as described previously (17). The purity of the SP-A preparation was confirmed by SDS-PAGE followed by silver staining and then filtered through a 0.2-micron filter to remove potential contaminants.

### Cells and Viruses

HEK293T-ACE2+TMPRSS2 (human embryonic kidney cell line overexpressing both human ACE2 and TMPRSS2 genes) and A549-ACE2 (human lung carcinoma epithelial cell overexpressing human ACE2, BEI Resources, NIAID) and Vero E6 cells (BEI resources, NIAID) were cultured and maintained in GlutaMax Dulbecco’s modified Eagle’s medium (DMEM) containing 1g/L D-glucose and 110mg/L sodium pyruvate supplemented with 10% (v/v) fetal bovine serum (FBS) and 1% (v/v) antibiotic (100U/mL of penicillin and 100µg/mL of streptomycin, Gibco) at 37°C and 5% CO_2_ atmosphere.

SARS-CoV-2 (Delta, B.1.617.2, 1.8×10^6^ PFU/ml, P3) used in this study was obtained from the World Reference Center for Arboviruses (WRCEVA) and was propagated in Vero E6 cells under BSL-3 containment conditions. Plaque forming assay (PFU) was performed to determine viral titer.

### Generation of hACE2 and TMPRSS2-Stably Expressing HEK-293T Cells

HEK293/ACE2/TMPRSS2 cell line, which is stable-producing human ACE2 and co-receptor, TMPRSS2, is a kind gift from Dr. Marc C Johnson of University of Missouri (18).

### Production of Luciferase- and GFP-Tagged Pseudotyped Particles

SARS-CoV-2 S protein gene expressing cDNA (gift from Dr. Marc Johnson, University of Missouri School of Medicine) was used to pseudotype Feline immunodeficiency virus (FIV) expressing luciferase (ScV2 S-FIV-mCherry/Luc) or green fluorescent protein (ScV2 S-FIV-GFP) by using previously described methods (18, 19).

### ELISA assays

Polystyrene microtiter plates were coated with 1 µg/ml recombinant SARS-CoV-2 spike protein produced in HEK293T cells (10549-CV-100, R&D Systems and Biotech, NE, MN, USA) while some plates were coated with 50 ng/ml SARS-CoV-2 S protein RBD or Biotinylated hACE2 (0.2 µg/ml) (EP-105; AcroBiosystems, Newark, USA) overnight at 4°C in sodium carbonate buffer (pH 9.6). The S protein has mutations that ensure its prefusion conformation. After coating, the plates were washed four times with TBST (pH 7.4-7.6) and blocked with 3% BSA diluted in TBS buffer for 1:30 mins at room temperature (RT). Then we added a series of two-fold dilutions of purified human SP-A (0-10 µg/ml, 100µl/well) in TBST containing either 5 mM CaCl_2_ (Sigma-Aldrich, Saint Louis, MO, USA) or 10 mM EDTA (Sigma-Aldrich) and incubated at room temperature for 1 h. The wells were washed 4 times and incubated with SP-A IgG polyclonal antibody (1:1000) at room temperature for 1 h with slight shaking and SP-A-S protein or SP-A-RBD complexes were detected by adding HRP-conjugated Goat Anti-rabbit IgG (1:2000, Bio-rad, Hercules, USA) for an additional 1 h. The absorbance (450 nm) of individual wells was quantified using a spectrophotometer (Multiscan Ascent, Labsystem; Fisher Scientific, NH, USA). Experiments were carried out in duplicates from three independent experiments.

### Competition Assays

SP-A (10 µg/ml) was incubated simultaneously in a buffer containing 10 mM each of either maltose, mannose, galactose, or N-acetylglucosamine for 1 h in plates previously coated with either S protein (1 µg/ml) or RBD (50 ng/ml, VANC00B, R&D Systems). As a control, SP-A was incubated in a 5 mM CaCl_2_-containing buffer without sugars. In another experiment, SP-A (10 µg/ml) was simultaneously incubated with 10 µg/ml biotinylated ACE2 in wells previously immobilized with SARS-CoV-2 RBD. Bound SP-A-S protein and/or RBD was detected with SP-A IgG polyclonal antibody (1:1000) and bound biotinylated ACE2 and RBD in the presence or absence of SP-A was detected by incubating with Streptavidin-HRP (VANC00B, R&D Systems) for 1 h at room temperature and developed as described above. All analyses were carried out in duplicate (n=3).

### Pseudotyped Virus Entry Assay in the Presence of human SP-A

HEK293T-ACE2+TMPRSS2 and A549-ACE2 cells were seeded in 24-well plates to obtain confluency after a 24 h culture. Luciferase- and GFP-tagged SARS-CoV-2 pseudotyped viral particles (MOI: 5; with wildtype (WT) S protein on their surface) were pre-incubated with varying concentrations of SP-A (0 to 50 µg/ml) in 1X MEM containing 5mM CaCl_2_ buffer for 1 h at RT. Cells were washed with 1X MEM and virus-SP-A mixture inoculated onto confluent cell monolayers and incubated at 37°C for another 2 h in a 5% CO_2_ incubator. Following this, the virus-protein mix was removed and fresh medium containing 2% FBS in DMEM (500 μl/well) was added to the cells and incubated at 37°C for another 48 h. To measure luciferase intensity in cells, the cells were washed gently with sterile PBS and lysed by incubating in cell culture lysis buffer for 15 mins at room temperature with slight shaking, and Firefly luciferase activity (RLU) was measured using Luciferase Assay System (E1500 kit, Promega, Madison, WI, USA). For GFP expression analysis, cells were mounted on coverslips and GFP intensity in the cells was observed by Nikon Eclipse TE2000-U microscope (Nikon Corporation, Tokyo, Japan), and GFP-expressing cells were quantified using the Image J software.

### SARS-CoV-2 (Delta Variant) Binding, Entry, and Infectivity Assays in the Presence of human SP-A

Binding assays was performed by pre-incubating infectious SARS-CoV-2 (B.1.617.2) with human SP-A for 1 h before inoculating pre-chilled A549-ACE2 cells with virus/protein mixture at 4°C for 2 h to allow virus binding to the cell surface. The cells were washed four times with cold PBS to remove unbound viral particles. Total RNA was isolated, and the amount of virus on cells was quantified by RT-qPCR. For entry assays, after 2 h incubation of SP-A + virus mixture at 4°C, the cells were washed, and fresh media was added and shifted to 37°C for 1 h to allow virus entry into cells. Then the cells were washed and treated with proteinase K (1 mg/ml) to remove attached viral particles on the cell surface and the amount of internalized viral particles was quantified by RT-qPCR. To further assess the role of SP-A in SARS-CoV-2 entry and infectivity, we pre-incubated virus with SP-A (0 to 50 μg/ml) for 1 h before inoculating confluent A549-ACE2 cells with the virus + protein mixture for 2 h at 37°C (MOI= 0.05), unbound virus was washed, and fresh growth media added, and cells incubated at 37°C and cells and supernatant collected at 4 h and 24 hpi and viral RNA, protein and titer in cells and supernatant analyzed.

### Plaque Assay

The potential antiviral activity of human SP-A against SARS-CoV-2 was detected by plaque assay in Vero E6 cells. Cell culture media were collected from A549-ACE2 cells inoculated with SP-A pretreated and untreated SARS-CoV-2 24 hpi and virus titer quantified as previously described(20). Briefly, Confluent monolayers of Vero E6 in 24-well plates were infected with 10-fold serial dilutions of supernatants from each treatment group at the indicated concentrations of SP-A (see results). The cells were cultured for 1 h with intermittent rocking. The unbound virus was removed and overlayed with 2% methylcellulose and cultured for another 72 h. Upon the development of plaques, cells were fixed with 10% formalin for 1 h and stained with 0.05% (w/v) crystal violet in 20% methanol and plaques were counted.

### Immunoblotting Analysis

Cells were lysed with RIPA buffer (Thermo Scientific, Rockford, IL) containing a cocktail of protease and phosphatase inhibitors (Roche, Indianapolis, IN). Total protein in cell lysates obtained at 4 hpi (to assess SP-A’s role in SARS-CoV-2 entry) and 24 hpi (to elucidate the effect of SP-A on viral infectivity) was determined using the BCA protein assay kit (Thermo Scientific). Five micrograms of total protein were resolved by SDS-PAGE on a 10% gel under reducing conditions and transferred to PVDF membranes (Bio-Rad). The blots were blocked in TBS containing 5% non-fat milk for 30 mins and incubated with SARS-CoV-2 nucleocapsid protein antibody (1:1000, NB100-56576, Novus Biological, CO, USA) overnight at 4°C. As a loading control, blots were stripped and re-probed with β-actin (1:1000, ab-16039, Abcam, MA, USA). Subsequently, the membranes were incubated with goat anti-rabbit IgG HRP-conjugated secondary antibody (Bio-Rad) and developed using ECL Western Blotting Substrate (Thermo Scientific).

### Real-time quantitative PCR (real-time qPCR)

Total RNA in cell lysates was extracted using a Quick-RNA extraction miniprep kit (# R1055 Zymo Research, CA, USA) following the manufacturer’s instructions and RNA concentration determined by the nanodrop machine (Thermo Scientific). Real-time RT-qPCR was performed using the AB StepOnePlus Detection System and the one-step kit RT-PCR Master Mix Reagents (#64471423, Biorad). Reaction mixtures were prepared according to the manufacturer’s instructions. In brief, 30 ng of RNA (we used 50ng in the binding and entry assays) was added into a 25 µl total volume of real-time PCR mix buffer containing forward/reverse primer pairs (forward, AGCCTCTTCTCGTTCCTCATCAC; reverse, CCGCCATTGCCAGCCATTC; each 500 nM) targeting SARS-CoV-2 N1 gene and a probe (250 nM, FAM) and other reagents provided by the manufacturer. The one-step q-RTPCR was carried out through one cycle of reverse transcription at 55°C for 10 mins followed by 40 cycles of amplification at 95°C for 3 mins, 95°C for 15 s, and 55°C for 1 min. The data were analyzed as fold change in CT values compared to SP-A untreated samples using the 2^-ΔΔCt^ method.

### Determination of Total Protein and SP-A Levels in Human Saliva Specimens

Saliva samples from 40 hospitalized COVID-19 patients and 12 healthy individuals were collected following IRB approval and SP-A level was assessed by ELISA. First, the total protein concentration of the individual saliva samples was analyzed. Following this, 5 µg/ml of individual saliva samples and purified human SP-A as a standard (0 to 0.05 µg/ml) were coated overnight on microtiter wells. Subsequently, the SP-A level was determined by measuring the absorbance as described above. All analyses were carried out in three independent experiments.

### Statistical analysis

All experimental data are presented as mean ± standard error and statistically analyzed using GraphPad Prism 8.0 (GraphPad Software, San Diego, CA, USA). Comparisons between two independent groups were performed using Student’s *t*-test or multiple groups using one-way ANOVA. It was considered statistically significant when P<0.05.

## Results

### SARS-CoV-2 S Protein and RBD are Recognized by Human SP-A protein

SP-A activates host innate immunity by binding to pathogen-associated molecular patterns (PAMPs) such as glycan units on viral surface proteins. Given the profound level of glycosylation on SARS-CoV-2 S-protein (5), and SP-A binding to other respiratory viruses (10, 21); we examined the potential interaction of human SP-A with SARS-CoV-2 S protein and the RBD. As shown in Figure 1A, SP-A bound to SARS-CoV-2 S protein in a dose-dependent manner (0 -10 μg/ml of SP-A) in the presence of calcium (5mM) but the binding of SP-A to S protein was reduced by 46% in the presence of 10 mM EDTA (a calcium ion chelator), suggesting that SP-A binding to S protein is slightly calcium-dependent (CRD domain) but other non-calcium-dependent regions might also play a role in S protein interaction because EDTA could not completely abrogate the SP-A binding to S protein. Similarly, a dose-dependent binding of SP-A (0 – 10 μg/ml of SP-A) to immobilized SARS-CoV-2 RBD was also observed (Figure 1B). As shown in Figure 1B, 59% decrease in SP-A binding to the RBD was observed in the presence of 10 mM EDTA, suggesting the involvement of SP-A CRD and other domains in the interaction between SP-A and RBD.

**Figure 1.**
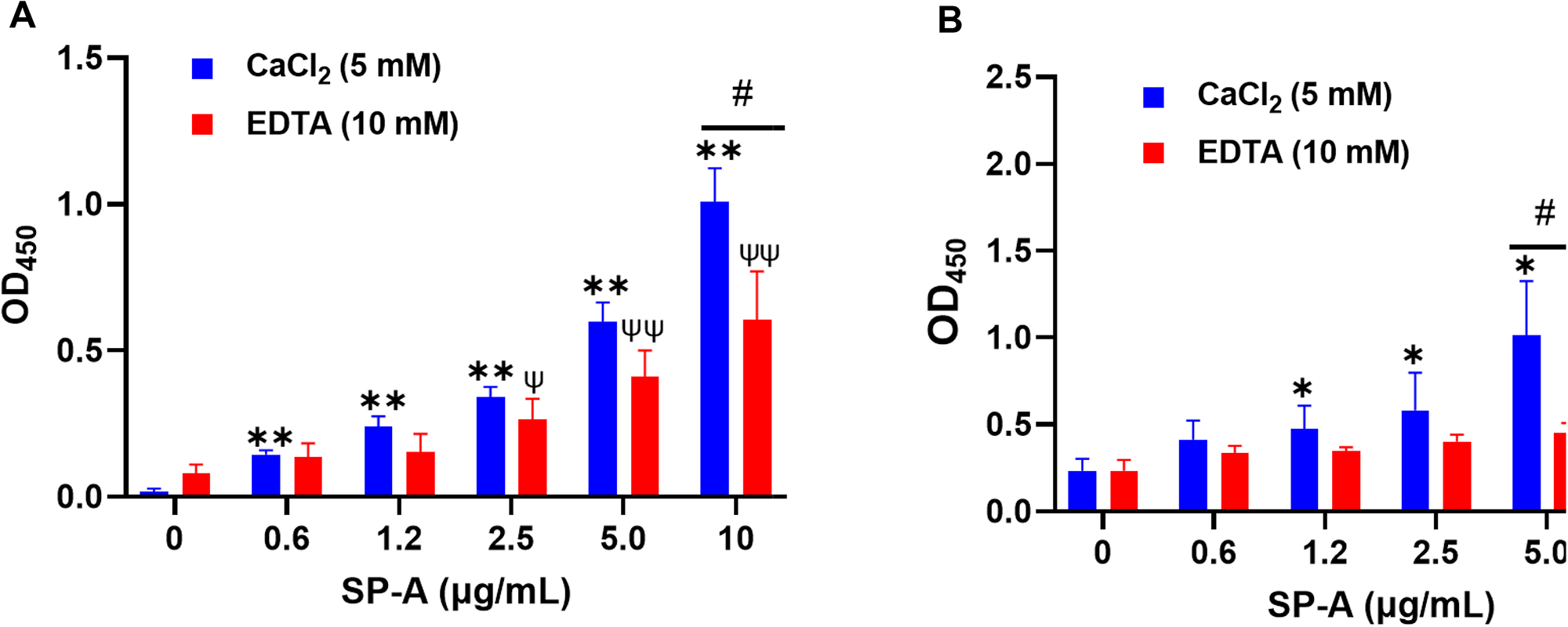
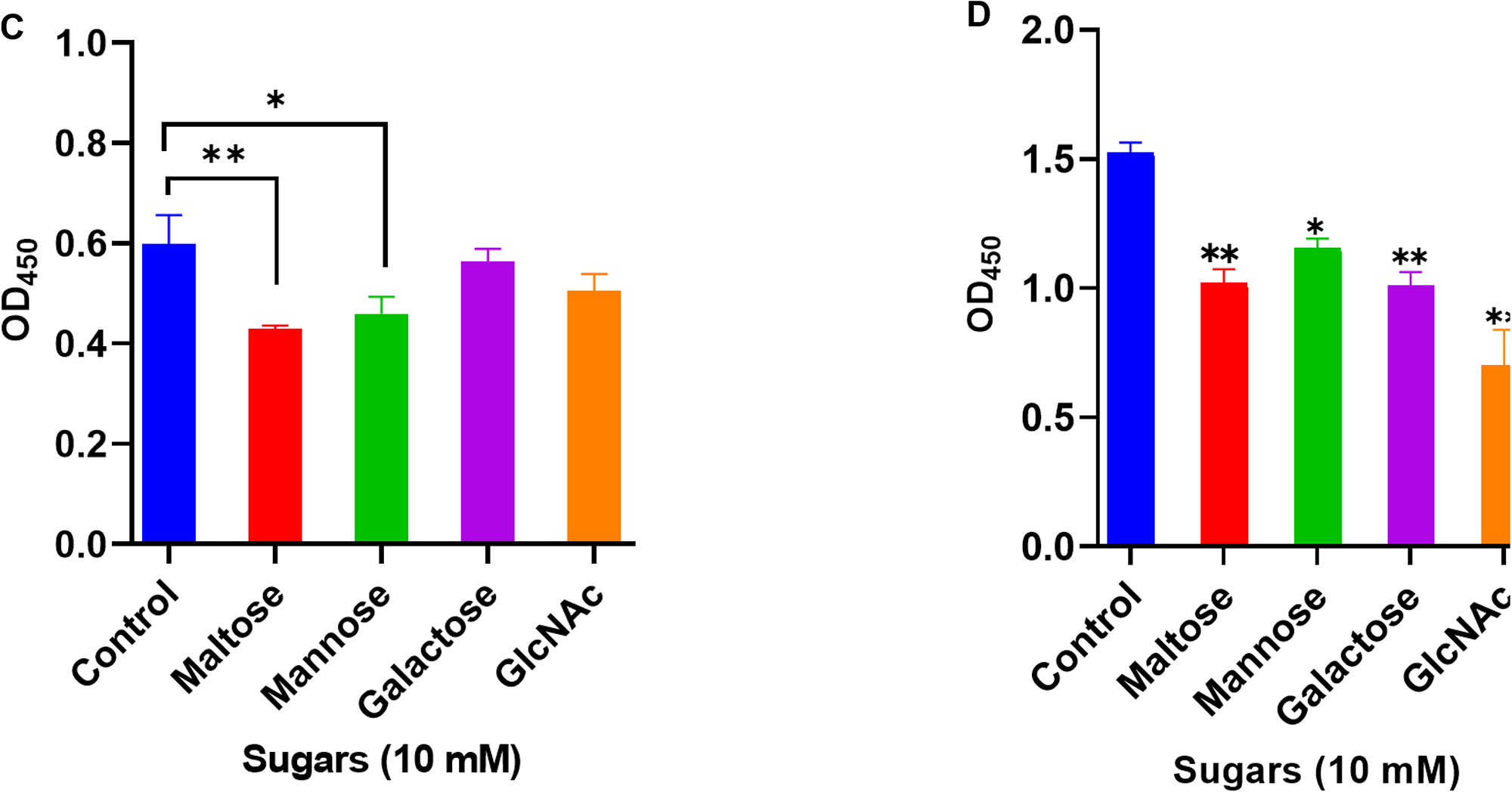
Human SP-A interacts with SARS-CoV-2 S protein and RBD. Purified S protein and RBD was immobilized on ELISA plates followed by incubation with serial dilutions of SP-A (0-10 μg/ml) in either 5 mM CaCl_2_ or 10 mM EDTA-containing buffer. The data show a dose-dependent increase in SP-A binding to SARS-CoV-2 S protein (A) and RBD (B). SARS-CoV-2 S protein (C) and RBD (D) coated plates were incubated with 10 μg/ml SP-A in the presence of 10 mM of each kind of sugar i.e. maltose, mannose, galactose, and N-acetylglucosamine (GlcNAc). Control samples were incubated with SP-A in 5 mM CaCl_2_ buffer without sugars and absorbance readings compared to control samples. Experiments were carried out in duplicates and three independent experiments. The data are presented as mean ± SE of 3 independent experiments. (*P<0.05; **P<0.01 reflect the levels of statistical significance in comparison with no SP-A treated group (0 μg/ml) in 5 mM CaCl_2_-containing buffer by unpaired student’s t-test analysis. ψ<0.05, ψψ<0.01 are the levels of significance compared to 0 μg/ml SP-A in 10 mM EDTA buffer while #<0.05 is the level of significance for the same SP-A concentrations in 5 mM CaCl_2_ vs 10 mM EDTA-containing groups.

### The binding of SP-A to SARS-CoV-2 S Protein and RBD is Inhibited by Sugars

Each monomer of SARS-CoV-2 S protein has about 17 of its 22 N-glycosylation sites occupied by glycans with two O-linked glycan sites on the RBD (6). SP-A has various binding affinities for carbohydrate molecules. Thus, we hypothesized that SP-A binds to S protein and RBD by recognizing the sugars displayed on its surface. To demonstrate whether sugars can competitively inhibit SP-A recognition of S protein and RBD, we incubated SP-A with immobilized S protein and RBD in the presence of 10 mM sugars (maltose, mannose, galactose, and GlcNAc). In Figure 1C, a reduced SP-A binding to S protein was observed in the presence of maltose and mannose with no significant inhibition by Galactose and GlcNAc. However, SP-A recognition of RBD was inhibited by all the sugars tested (Figure 1D). Although, increasing concentrations of maltose could not completely abrogate SP-A interaction with S protein and RBD (Supplementary Figure E1 A&B). These data indicate that SP-A binds glycoconjugates on the S protein and RBD of SARS-CoV-2. However, there is also the potential for other forms of protein-protein interaction because of the observed interaction regardless of the presence or dose of sugars.

### SP-A Binds to hACE2 and Impacts SARS-CoV-2 RBD Interaction with hACE2 Receptor

Since SARS-CoV-2 mainly infects hACE2-expressing airway and lung epithelial cells which are also the predominant SP-A-expressing cells; we assessed whether SP-A could interact with hACE2 directly. We observed a dose-dependent increase in SP-A binding to biotinylated hACE2 (Figure 2A). Furthermore, SP-A binding to hACE2 decreased significantly in the presence of EDTA, indicative of its reliance on calcium for optimum interaction with hACE2. However, Mannose, a preferred ligand for SP-A, showed weak inhibitory effect to SP-A interaction with hACE2, suggesting that SP-A interaction with hACE2 may act through glycoconjugate and non-glycoconjugate binding (Suppl. Figure E2). Given that SP-A can bind hACE2, we further examined its impact on the interaction between SARS-CoV-2 RBD and hACE2 by simultaneously incubating SP-A and hACE2 on ELISA plates immobilized with RBD. We observed a reduced hACE2 binding to RBD in the presence of SP-A while BSA (used as a control for non-specific protein interaction) had no inhibitory effect on RBD-hACE2 interaction (Figure 2B). Therefore, SP-A can interfere with the interaction between RBD and hACE2.

**Figure 2.**
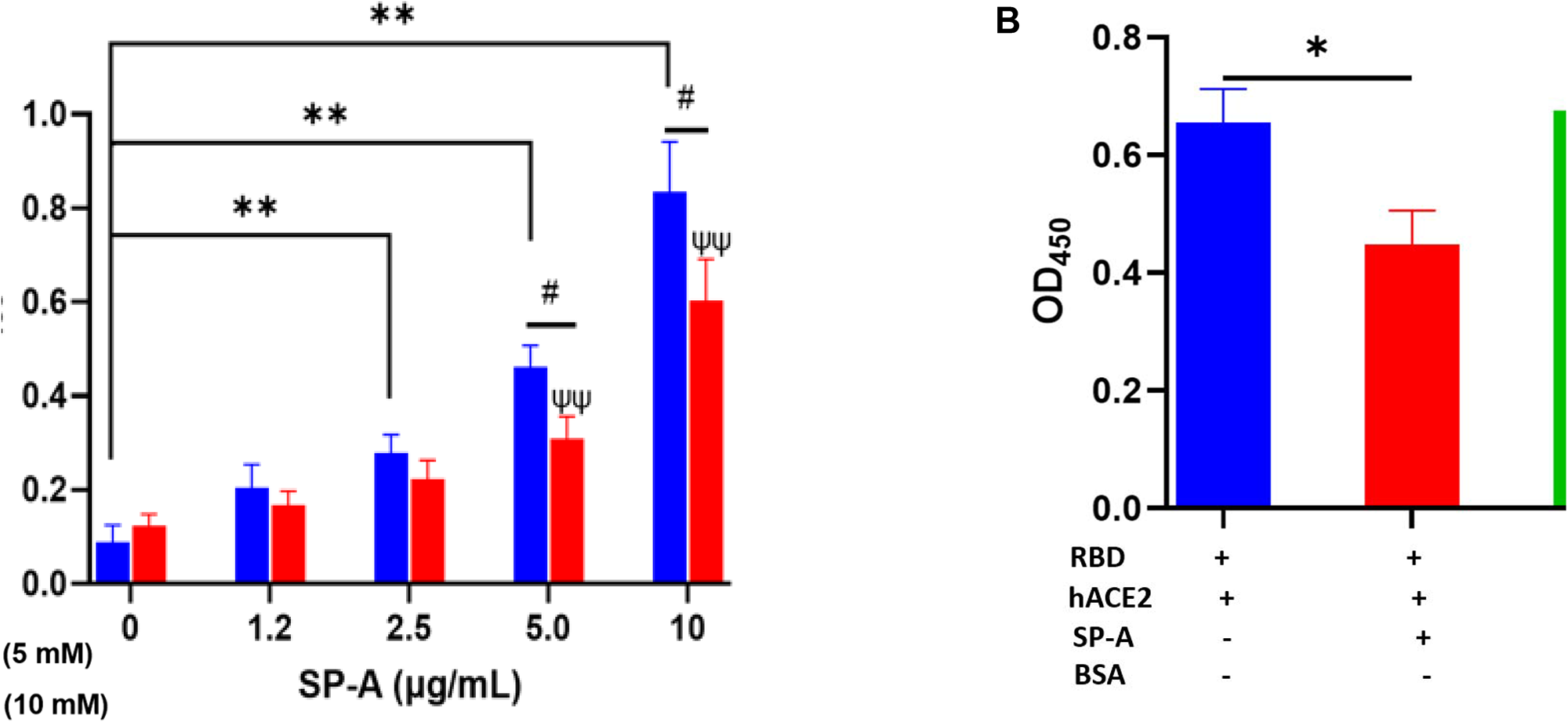
SP-A binds to hACE2 and impacts the interaction of SARS-CoV-2 RBD and hACE2. (A): Microtiter plates were coated with biotinylated hACE2 (0.2 μg/ml) and plates were then incubated with a range of SP-A concentrations (0-10 μg/ml) in the presence of 5 mM calcium or 10 mM EDTA, and the level of bound SP-A was detected as described above. (B): SP-A (10 µg/ml) was simultaneously incubated with biotinylated hACE2 (10 µg/ml) in wells immobilized with RBD (5 µg/ml). To assess non-specific protein interaction, some wells were incubated with 10 µg/ml BSA. Statistical analysis was performed by a student’s t-test. Values are mean ± SEM (*P<0.05, **P<0.01 compared to 0 μg/ml SP-A in CaCl_2_-containing group. ψψ<0.01 vs 0 μg/ml SP-A in 10 mM EDTA group while #<0.05 is the level of significance for same SP-A concentrations in 5 mM CaCl_2_ vs 10 mM EDTA-containing groups.

### Human SP-A Inhibits Binding and Entry of SARS-CoV-2 Pseudotyped Particles and SARS-CoV-2 (Delta variant) in host Cells

The ELISA showed that SP-A binds to SARS-CoV-2 S protein and RBD, as well as interfere with RBD-hACE2 interaction. We further examined whether SP-A could inhibit SARS-CoV-2 infectivity using both pseudotyped particles and infectious SARS-CoV-2 (Delta variant). It is known that glycans adjacent to the RBD can serve as determinants of viral binding with cellular receptors to facilitate subsequent entry (22), and SP-A interaction with RBD was attenuated in the presence of sugars which suggests that SARS-CoV-2 entry in hACE2-expressing host cells may be inhibited by pre-treating with SP-A. Thus, we assessed whether the capacity of SP-A to bind S protein and RBD can result in viral entry inhibition. We challenged HEK293T-ACE2+TMPRSS2 and A549-ACE2 cell lines with a luciferase- or GFP-tagged SARS-CoV-2 pseudotyped particle (expressing wildtype (WT) S protein on its surface) pre-incubated with or without SP-A and observed significantly reduced luciferase and GFP intensities in an SP-A dose-dependent manner starting at 6.25 μg/ml in HEK293-ACE2+ TMPRSS2 (Figure 3A) and 12.5 μg/ml in A549-ACE2 (Figure 3B). As shown in Figures 3C & 3D, we also observed a decrease in GFP intensity with increasing SP-A concentrations in a dose-dependent manner in HEK-293T-ACE2 cells.

**Figure 3.**
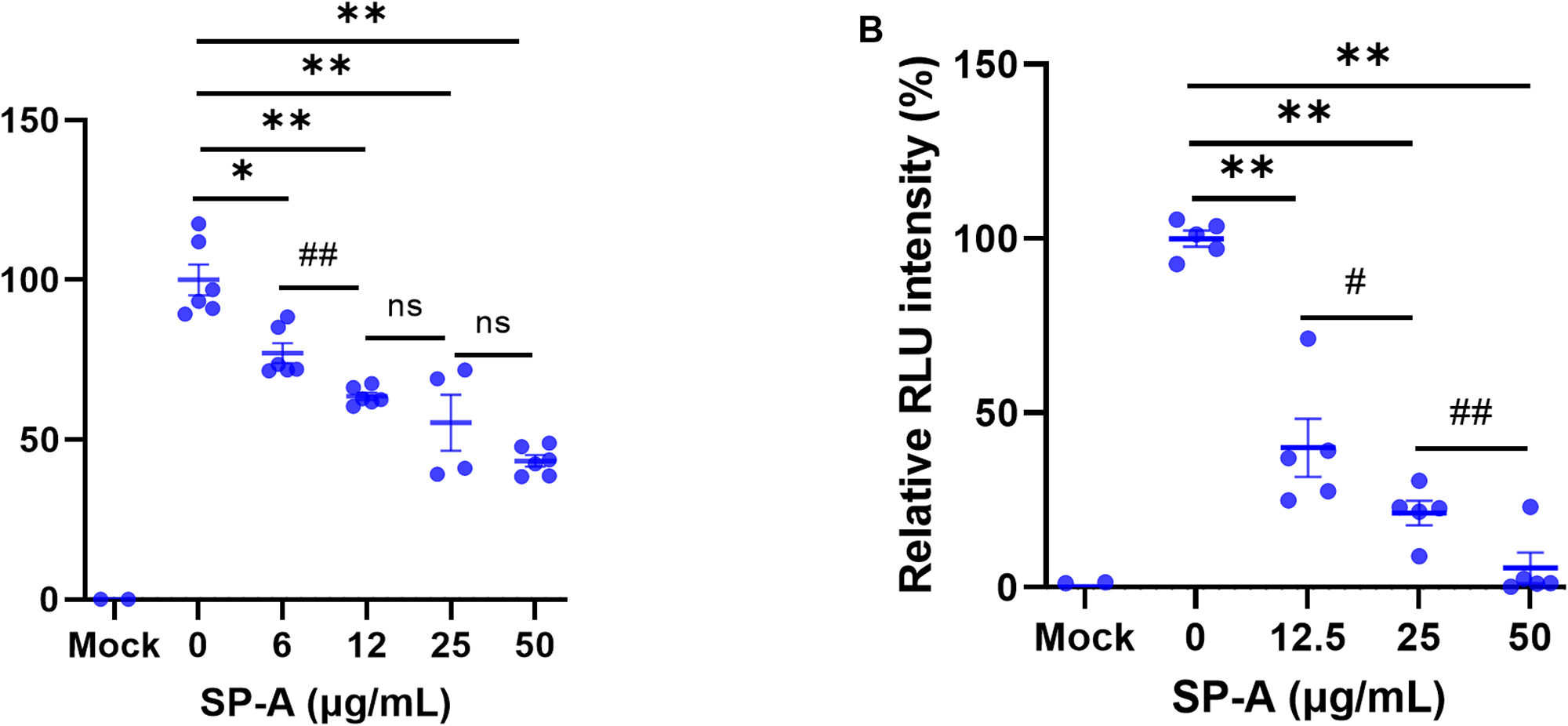
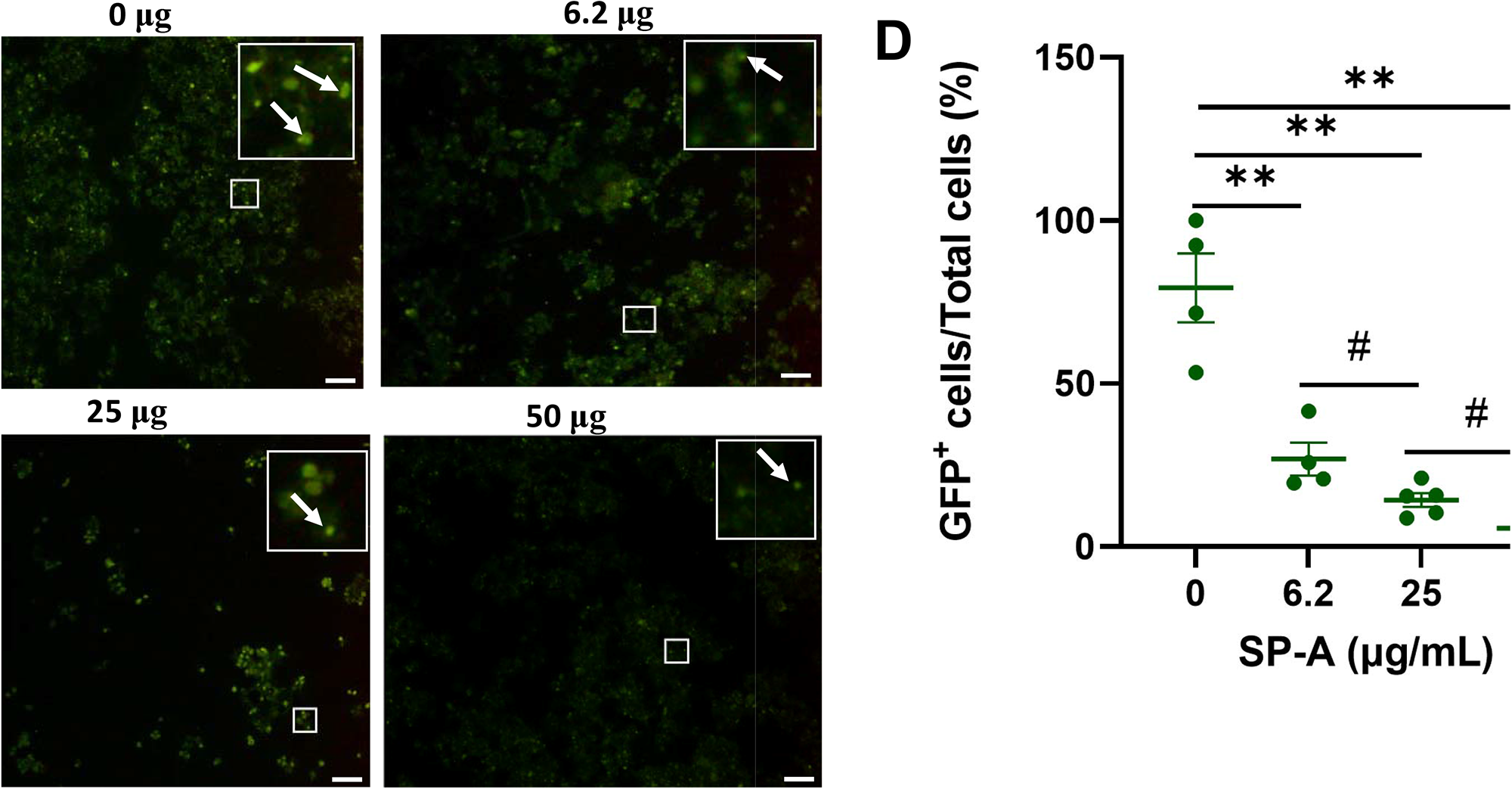
SP-A inhibits entry of SARS-CoV-2 pseudotyped lentiviral particles in susceptible host cells. We measured the luciferase intensity of the SP-A (0-50 μg/ml) pre-treated WT S protein pseudotyped lentiviral particles in the cell lines: (A): HEK293T-ACE2+TMPRSS2; (B): A549-ACE2. (C): Representative images of GFP-positive cells in HEK293T-ACE2+TMPRSS2 challenged with SP-A pre-treated and un-treated GFP-tagged SARS-CoV-2 pseudotyped particles. (D): GFP signal taken from 10 images per well, normalized to the cellular area and quantified by Image J software. The data were expressed as mean ± SEM of the percentage of positive cells/total cells per area analyzed (n=3). Scale bar = 100 μm. *P<0.05; **P<0.01, compared to control sample (0 μg/ml).

To further validate the role of SP-A in SARS-CoV-2 infectivity, binding, and entry assays with an infectious SARS-CoV-2 (B.1.617.2, Delta variant) were performed. We observed that the level of viral particles bound on the cell surface decreased significantly in a dose-dependent manner compared to the BSA-treated and untreated (0 μg/ml) controls at 4°C (Figure 4A). Temperature shift experiments at 37°C for 1 h showed lower levels of internalized viral RNA with increasing SP-A concentrations (Figure 4B). This level of the internalized virus was reduced by ≈ 58% relative to control (0 μg/ml) in cells inoculated with virus + 25 μg/ml SP-A. Before shifting to 37°C, we validated viral entry by treating some cells inoculated with virus only (0 μg/ml SP-A) with proteinase K to remove virus on the cell surface after the 4°C incubation (binding control) and observed only minimal levels of bound viruses. As previously described with pseudotyped viruses, pre-incubation of virus with increasing SP-A concentrations (0 to 50 μg/ml) before inoculating confluent A549-ACE2 cells resulted in a dose-dependent decrease in SARS-CoV-2 N protein levels in cells 4 hpi (Figures 4C and 4D). As expected, Chloroquine (CQ), an established inhibitor of SARS-CoV-2 entry used as a positive control, inhibited virus entry. The results highlight the binding and entry inhibitory activity of SP-A in the context of SARS-CoV-2 infection.

**Figure 4.**
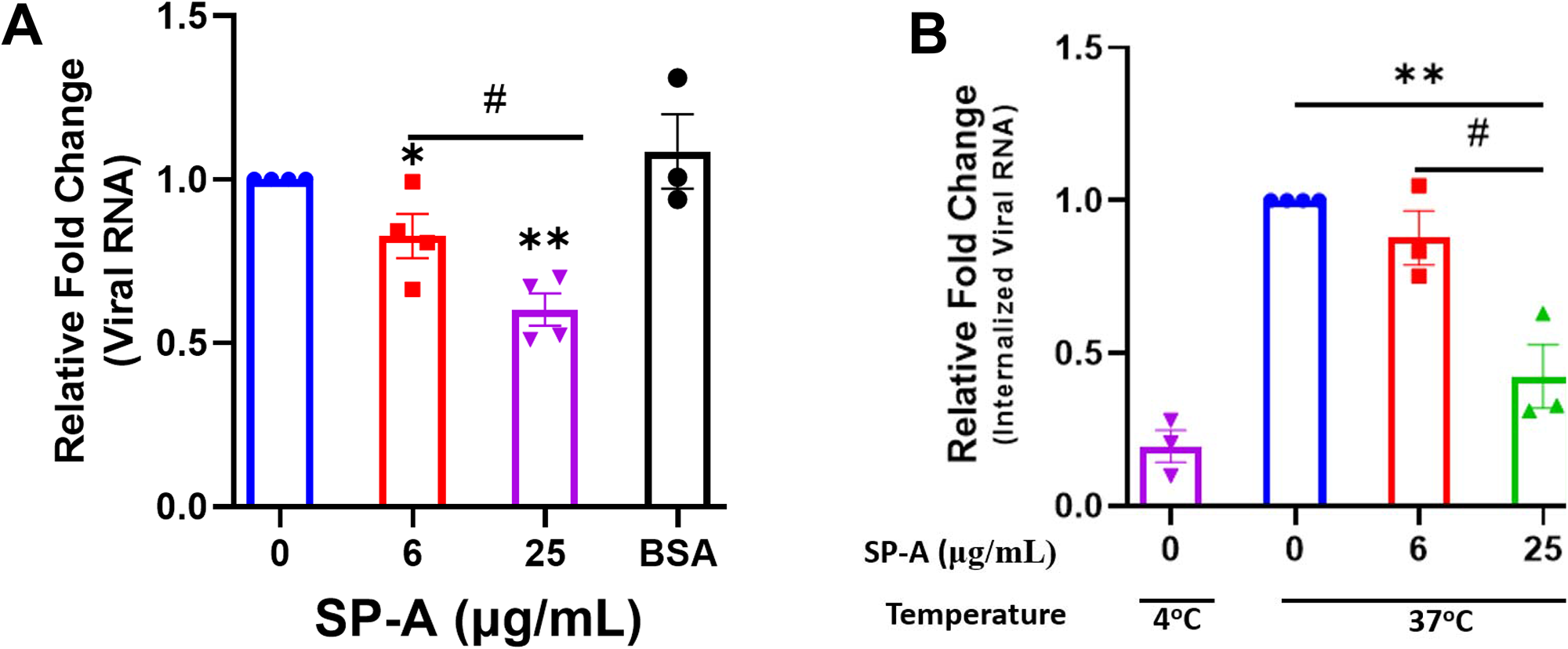
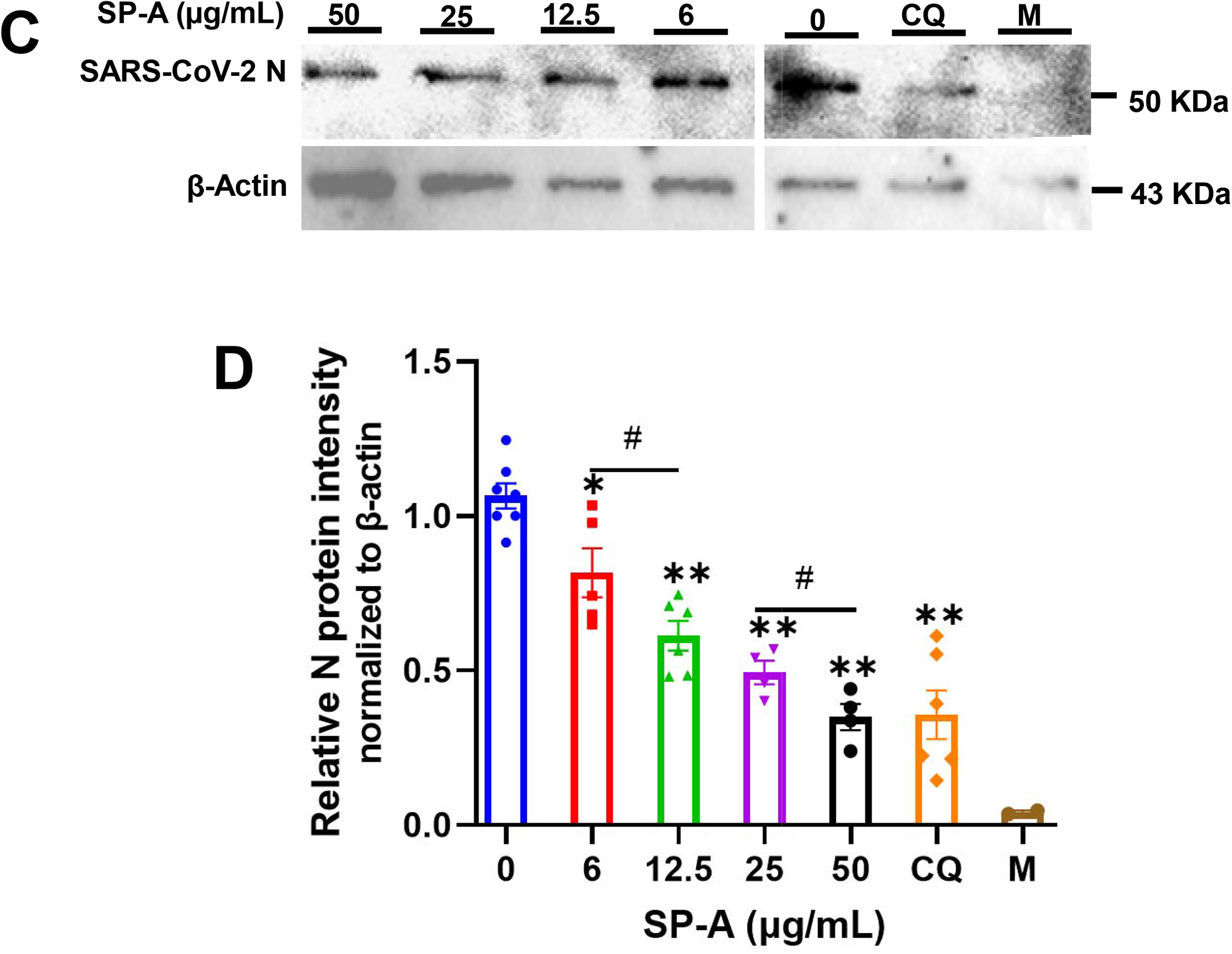
SP-A attenuates SARS-CoV-2 (Delta) binding and entry in A549-ACE2 cells. (A) Viral binding assays were performed in A549-ACE2 cells. SARS-CoV-2 (Delta variant) was pre-incubated with the indicated concentrations of SP-A or BSA (used as a non-specific protein control, 50 μg/ml) for 1h at RT. Then inoculated onto pre-chilled cells for another 2 h at 4°C to allow binding to the cell surface. (B): Viral entry assays were performed as described above. However, after 2 h incubation of SP-A + virus mixture at 4°C, the cells were washed, and fresh media was added and shifted to 37°C for 1 h to allow virus entry into cells. The cells were washed and treated with proteinase K (1 mg/ml) to remove attached viral particles on the cell surface and the amount of internalized viral particles was quantified by RT-qPCR. Binding control at 4°C was also used to assess virus entry by treating cells inoculated with virus only (0 μg/ml SP-A) after 2 h incubation with proteinase K prior to shifting to 37°C. The relative fold change was normalized to 18S rRNA internal control and expressed as mean ± SEM of the relative fold change in CT values compared to the control sample (0 μg/ml). (C): A549-ACE2 cells inoculated with SP-A + virus mixture for 2h and then cells were harvested after 4 h incubation for Western blotting analysis using SARS-CoV-2 N protein and β-actin antibodies, respectively. Representative images of Western blotting analysis of cell lysates with SARS-CoV-2 N protein and β-actin as a control. Chloroquine (CQ) (10 µM) was used as a positive control. (D): Quantification of N protein level relative to β-actin (loading control). Each data represents the mean (%) ± S.E. *P<0.05; **P<0.01, compared to control group (0 μg/ml); #<0.05= significance between two doses.

### Human SP-A Attenuates SARS-CoV-2 Infectivity in A549-ACE2 Cells

The hallmark of viral infectivity is the ability to not merely get into a cell but to also replicate its genome, express both structural and non-structural proteins, and assemble these components to produce infectious progeny viral particles. We therefore investigated the potential inhibitory role of SP-A in SARS-CoV-2 infectivity by RT-qPCR, immunoblotting and plaque assay using total RNA and protein isolated from infected cells and supernatant 24 h after infection of A549-ACE2 cells. The results showed that SP-A significantly reduced SARS-CoV-2 RNA level in cells in a dose-dependent manner (0 – 50 µg/ml of SP-A) (Figure 5A). In the presence of 50 µg/ml SP-A, we observed approximately 50% decrease in viral RNA level in cells. These results were further confirmed by immunoblotting assay where a dose-dependent decrease in the level of SARS-CoV-2 N protein was observed (Figure 5B & 5C). 50 µg/ml SP-A resulted in an approximately 10-fold reduction in virus titer compared to cells infected with the virus only by plaque assay (Figure 5D-E). Taken together, these results suggest that human SP-A has an inhibitory effect on SARS-CoV-2 entry and reduces viral loads in human lung epithelial cells by interacting with SARS-CoV-2 S protein and preventing viral binding to A549-ACE2 cells.

**Figure 5.**
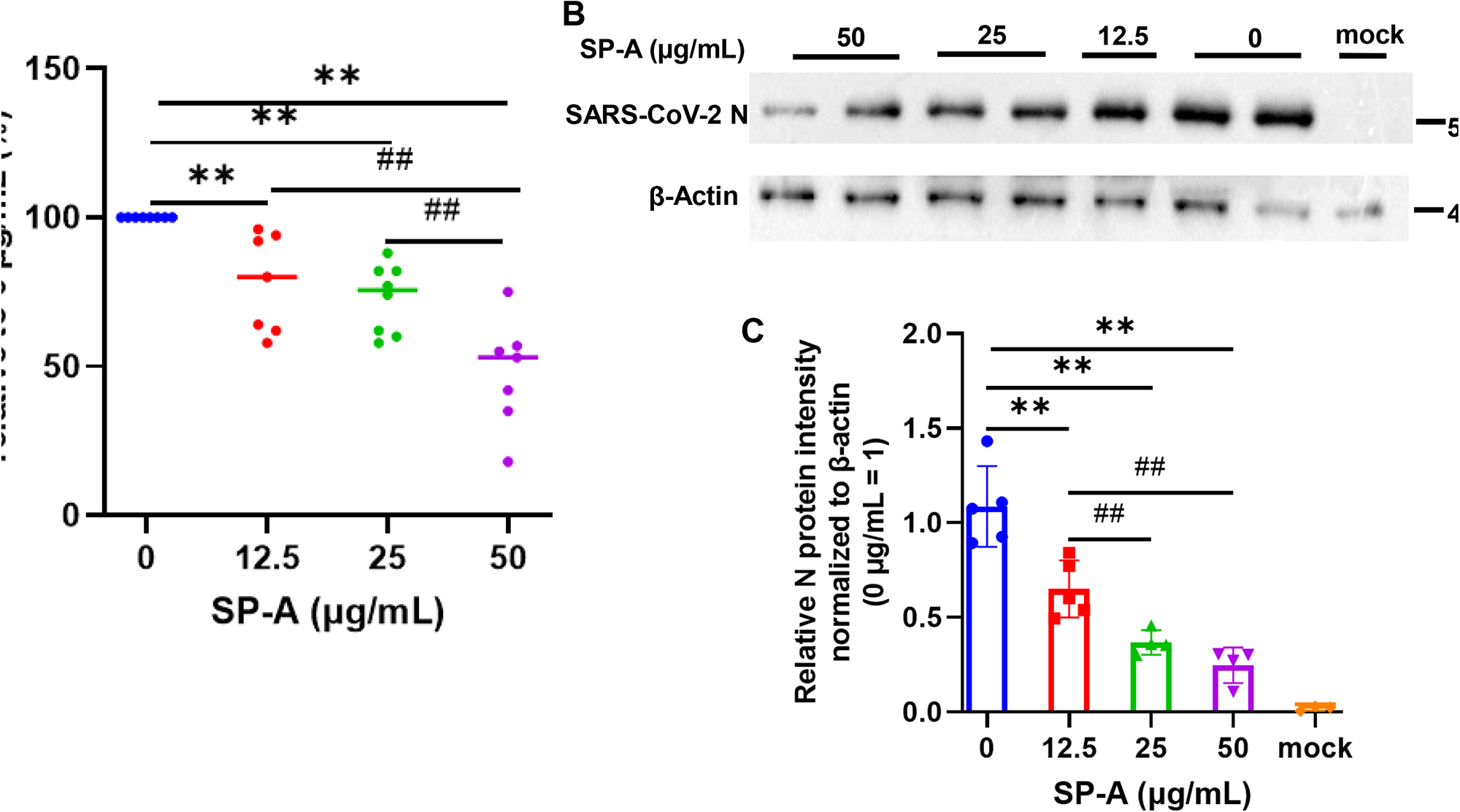
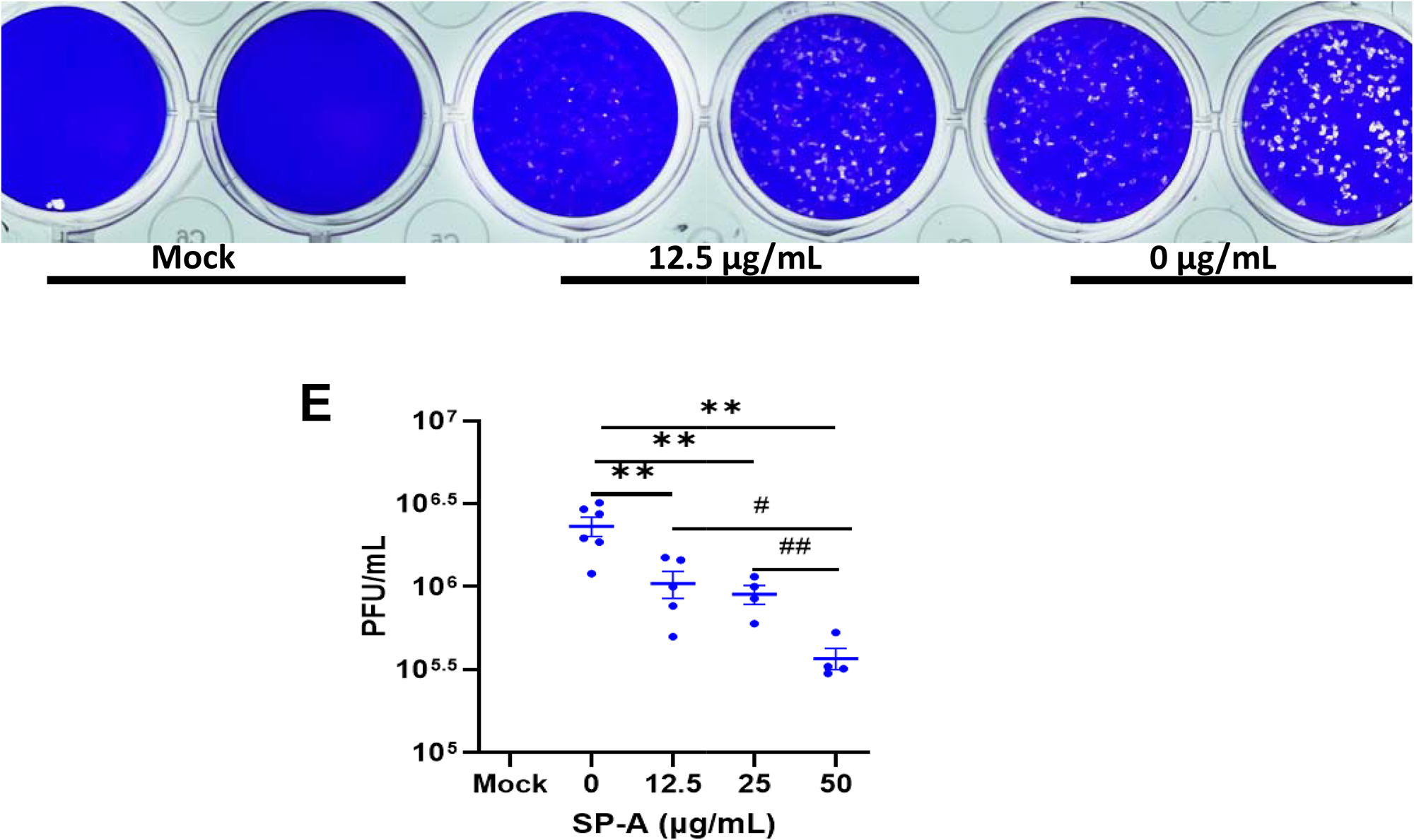
SP-A Inhibits SARS-CoV-2 (Delta) infectivity in A549-ACE2. SARS-CoV-2 was pre-incubated with increasing concentrations of SP-A before inoculating A549-ACE2. Cell culture media and cells were collected and used for RT-qPCR, western blot, and plaque assays. (A): Significant decrease in viral RNA levels in A549-ACE2 cells treated with or without SP-A and expressed as fold change in CT values relative to control (0 μg/ml) 24h hpi. (B): Western blots of SARS-CoV-2 N expressions in cell lysates treated with increasing concentrations of SP-A (C): Relative viral N protein levels in A549-ACE2 cell lysate normalized to β-actin. (D): Visible plaques on Vero E6 cells infected with or without (0 μg/ml) SP-A. (E): Dose-dependent decrease in virus titer with increasing SP-A concentrations. Values represent mean ± S.E. *P<0.05, **P<0.01, compared to the control sample (0 μg/ml) and #<0.05, ##<0.01 when compared between two concentrations (n= 3)

### Increased SP-A Level in the Saliva of COVID-19 Patients Compared to Healthy Controls

Changes in pulmonary surfactant and surfactant protein levels and elevated mannose-binding lectin (a related collectin in innate immunity) in the sera of some SARS and COVID-19 patients have been shown to correlate with disease severity (23–25). Thus, we examined the level of SP-A in the saliva of COVID-19 patients hospitalized with varying disease severity. The results showed that COVID-19 patients have an elevated total protein and SP-A levels in their saliva compared to healthy controls (Figure 6C). However, upon stratification based on disease severity, we observed a profoundly reduced SP-A level in the subgroup of severe COVID-19 patients (Figure 6D) despite the higher total protein levels observed in this subgroup (Figure 6B); highlighting the importance of a relatively preserved level of SP-A in the salivary mucosa of SARS-CoV-2 infected individuals to alleviate the most severe COVID-19 symptomology.

**Figure 6.**
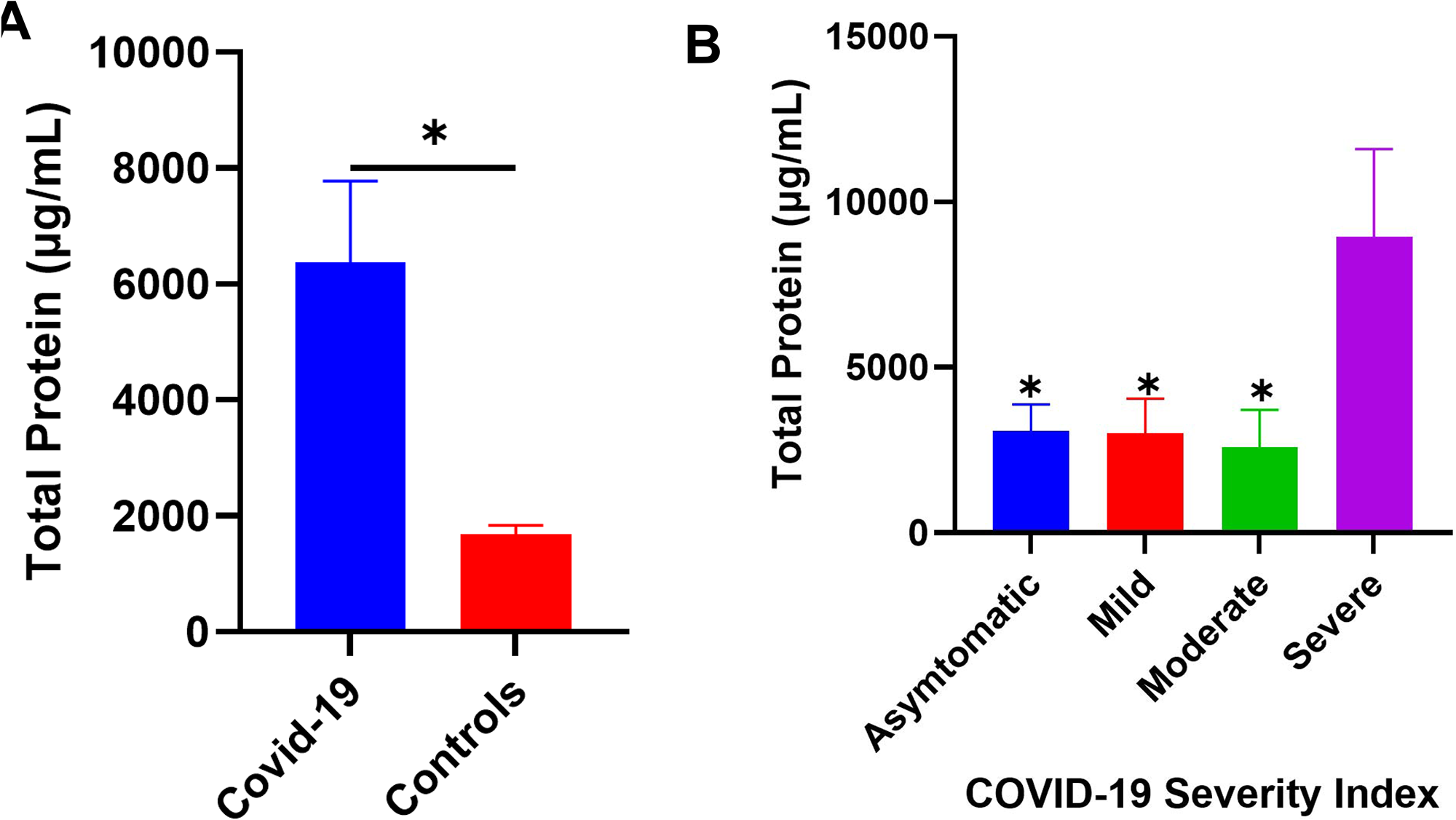
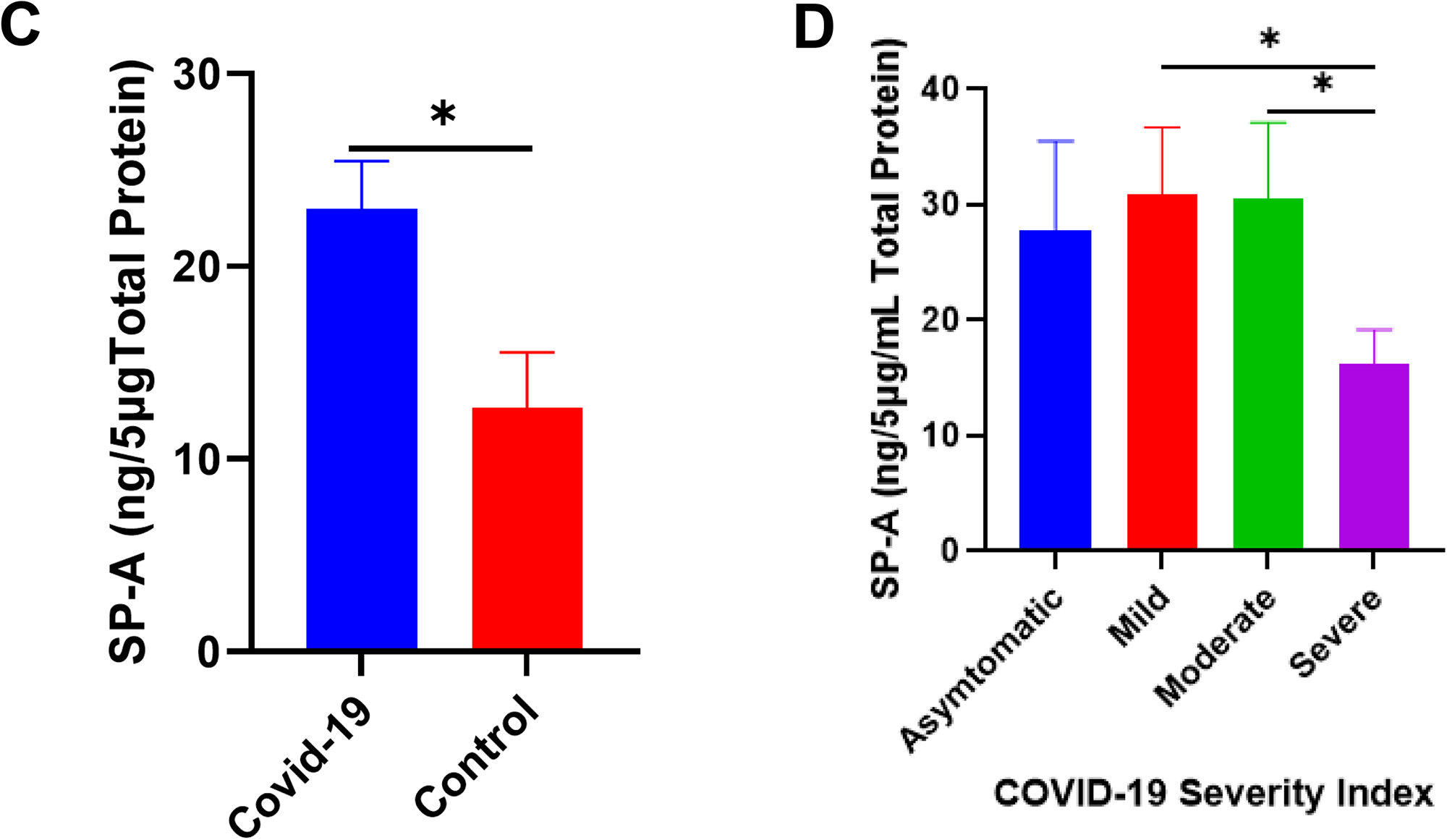
Increased SP-A level in the saliva of COVID-19 patients compared to healthy control. (A): Higher total protein concentration in the saliva of COVID-19 patients (n=40) and healthy controls (n=12). (B): Severe COVID-19 patients have significantly elevated total protein compared to asymptomatic, mild, and moderately infected patients. (C): Higher SP-A levels in the saliva of COVID-19 patients compared with healthy controls. (D): Upon stratification, severe patients had significantly reduced SP-A levels compared to mild and moderate patients.

## Discussion

Barely three years since first reported, COVID-19 has become one of the leading causes of death in the US (1). The goal of current antiviral research is to develop novel therapies that not only target viral proteins but also important host proteins/pathways that are essential for the virus’s life cycle (16, 26). As the current COVID-19 pandemic has highlighted with the frequent emergence of several variants with potential for immune escape, there is the need to rather focus on host antiviral proteins (16, 26). Moreover, the potential of using lectins (carbohydrate-binding proteins) has gained huge appeal in recent times since these molecules interact with relatively conserved glycoconjugates on viral proteins to mediate their antimicrobial functions (27). Thus, lectins are less sensitive to loss of function mutations due to sequence changes in viral surface proteins required for cellular entry and replication. Interestingly, SP-D was recently shown to inhibit SARS-CoV-2 entry and replication in the host cells (28). Therefore, we hypothesized that SP-A can bind to SARS-CoV-2 S protein and RBD and this interaction will influence viral binding, entry, and infectivity in susceptible host cells.

For the first time, our findings demonstrate that human SP-A can bind SARS-CoV-2 S protein and RBD in a dose-dependent manner. The biological significance of SP-A binding to SARS-CoV-2 S protein was emphasized by its inhibitory effect on viral infectivity in a lung epithelial cell line. Compared to RBD, we observed that SP-A interaction with the S protein is less dependent on calcium ions. The significantly reduced SP-A binding to RBD observed in the presence of the calcium ion chelator, EDTA, and binding competitors (sugars), signifies the involvement of the CRD whose binding capacity to glycans is dependent on divalent ions like calcium. Notably, the SARS-CoV-2 RBD is a 223 peptide (amino acid sequence from 319 to 541 of S protein), which might have influenced the capacity of other regions of the bouquet-shaped SP-A to recognize it (5). Meanwhile, the calcium-independent binding to the S protein observed in our study is in line with previous reports of SP-A’s interactions with the glycoproteins of IAV and HIV (though at low pH) (11, 12). Benne and colleagues previously showed that SP-A interacts with sialic acids in the hemagglutinin of IAV through its collagen-like domain (11). Our finding is novel because SP-A only partially recognized the S protein of a closely related beta coronavirus, SARS-CoV-1.

Since hACE2-expressing cells are the major targets for SARS-CoV-2 infection, we tested whether SP-A can interact with hACE2 and the role of SP-A in RBD-hACE2 interaction. We observed a calcium-dependent binding of SP-A to hACE2. This could indicate that SP-A also binds to hACE2 through a CRD-dependent mechanism. Importantly, the observation that SP-A can competitively attenuate RBD-hACE2 interaction supports the idea that SP-A may interfere with SARS-CoV-2 interaction with the host hACE2 receptors.

We next assessed the biological significance of SP-A binding with SARS-CoV-2 S protein by pre-incubating pseudotyped and infectious SARS-CoV-2 (Delta) with SP-A before cellular challenge. The delta variant was used for this experiment and in our infectivity assays because of the profound pathogenesis induced by this variant both in the human population and in *in vitro* and *in vivo* models of infection (29–31). In examining the early events that occur upon SP-A interaction with SARS-CoV-2, we have uncovered that SP-A binding to SARS-CoV-2 RBD can impair viral binding and entry in lung epithelial cells, ultimately resulting in low viral load in cells. Besides the low levels of viruses bound to cells surface at 4°C in the presence of SP-A, we observed significantly reduced viral RNA in cells 1 h after infection at 37°C compared to SP-A untreated controls. At 1 hpi, whatever viral gene detected in cells is reasoned to have been introduced by the infecting particles and not a result of viral replication in the infected cells which was supported by the reduced viral N protein 4 hpi in A549-ACE2 cells challenged with SP-A pre-treated virus (32). Chloroquine, an established entry inhibitor of SARS-CoV-2 was used as a positive control for this assay. The inhibitory role of chloroquine is mediated by its capacity to raise endosomal pH to halt the low pH-induced fusion of viral and endosomal membranes (33). These findings strongly suggest that the interaction between SP-A and viral S protein may have obstructed viral attachment to hACE2 on susceptible host cells, inhibiting SARS-CoV-2 entry as was previously observed with SP-A binding to HIV gp120 to prevent its interaction with CD4^+^ cells (12).

However, beyond binding and entry, a viral infection is deemed successful following genome replication, expression of structural and non-structural proteins, packaging of viral components, and release of infectious progeny viral particles. Our RT-qPCR, immunoblot, and plaque assays all demonstrate the inhibitory effect of SP-A on SARS-CoV-2 infectivity by the observed low levels of RNA, nucleocapsid protein and virus titer upon SP-A pre-treatment. The approximately 10-fold decrease in infectious virus titers and approximately 50% reduction in viral RNA at 50 µg/ml SP-A is quite outstanding and we speculate that the reduction in SARS-CoV-2 binding, and entry by SP-A resulted in the significantly low levels of virus load in permissive lung epithelial cells. This finding is supported by a recent study that SARS-CoV-2 infected clinical viral samples treated with SP-D had lower RNA-dependent RNA polymerase gene (RdRp); suggestive of the role of collectins as inhibitors of SARS-CoV-2 infectivity.

This study focused on pre-treating SARS-CoV-2 with SP-A before cellular challenge and the downstream effects on infectivity. However, a previous report showed that some lectins are more potent inhibitors of virus infection when cells are treated with lectins as against virus treatment before infection (34, 35). Since our ELISA assays showed that SP-A can recognize hACE2 receptors and interfere with RBD-hACE2 interaction, further mechanistic studies to characterize the antiviral effect of SP-A by pre-treating cells before SARS-CoV-2 challenge and the complexity of such an interaction merits further studies.

Interestingly, the same cells i.e. ATII cells that mainly produce and secrete collectins are also the predominant lung epithelial cells targeted by SARS-CoV-2 (2, 36). Infection and subsequent damage of these cells results in decreased SP-A production and secretion, rendering the lung more susceptible to injury (37–39). The importance of surfactant proteins has been demonstrated by the routine administration of surfactant-based replacement therapies in neonates with impaired lung functions (40). Thus, we assessed the levels of SP-A in the saliva of a subset of COVID-19 patients. Compared to controls, COVID-19 patients had relatively higher SP-A levels in their saliva, an increase that is suggestive of SP-A’s innate immune roles during an acute infection prior to the induction of the adaptive immune response. However, patients with severe disease had remarkably reduced SP-A levels compared to moderately and mildly infected patients. We thus speculate that basal levels of SP-A in salivary mucosal are increased upon SARS-CoV-2 infection; however, in severe patients, SP-A levels are remarkably depleted due to reduced expression or more degradation (41), thus impairing host innate antiviral response. A dysregulation in essential surfactant protein (SP) genes among COVID-19 patients has also been observed (42).

A limitation of our study is the small size of human saliva specimens used in this study, and that we do not know the time from infection to symptomology and hospitalization among our study participants and as a result cannot definitively link low SP-A levels to disease severity. For example, it could be reasoned that severe patients with low SP-A levels may have been hospitalized longer and may have had other pre-existing conditions and co-infections that could have seriously impacted SP-A levels in their saliva. Indeed, SARS-CoV-2 infection results in profound lymphocytopenia, that can predispose individuals to secondary infections by otherwise relatively nonpathogenic and pathogenic bacteria (43). In addition, there have been several reports of bacterial coinfections among very severe COVID-19 patients in the ICU (44, 45). The higher tendency for secondary bacterial infections among COVID-19 patients might be due to epithelial cell damage and/or alterations in host innate immune molecules and should be further investigated.

Several studies have previously shown that levels of collectins among the general population vary depending on an individual’s unique SP variant and this could result in differential SP-A levels (46). Structural and functional changes in surfactant proteins have been linked to single nucleotide polymorphisms demonstrated to affect their abilities to bind microbial PAMPs and carbohydrates (47). These polymorphisms in SP genes may influence their interactions with SARS-CoV-2 glycoproteins and future research should focus on elucidating the ability of SP-A variants to interact with SARS-CoV-2 glycoproteins and the functional roles of the variants on viral infectivity and pathogenesis both *in vitro and in vivo*. Moreover, since several SARS-CoV-2 variants have been observed with mutations in their S protein demonstrated to confer resistance to neutralization antibodies (14, 15, 48), it could mean that changes in S protein can remove or introduce novel N- and O-glycosylation sites that could potentially make the S protein more or less sensitive to SP-A as observed previously with IAV strains (6, 49). To address this possibility, further studies should examine mechanistically SP-A interactions with SARS-CoV-2 variants and the biological significance using both *in vitro* and animal models of infection.

In conclusion, our findings show that SP-A exerts strong antiviral effect against SARS-CoV-2 infectivity in lung epithelial cells by interacting with S protein, inhibiting viral binding, entry, and titer in susceptible cells (Figure 7). These findings supplement current efforts aimed at developing novel surfactant-based therapies to combat COVID-19.

**Figure 7.**
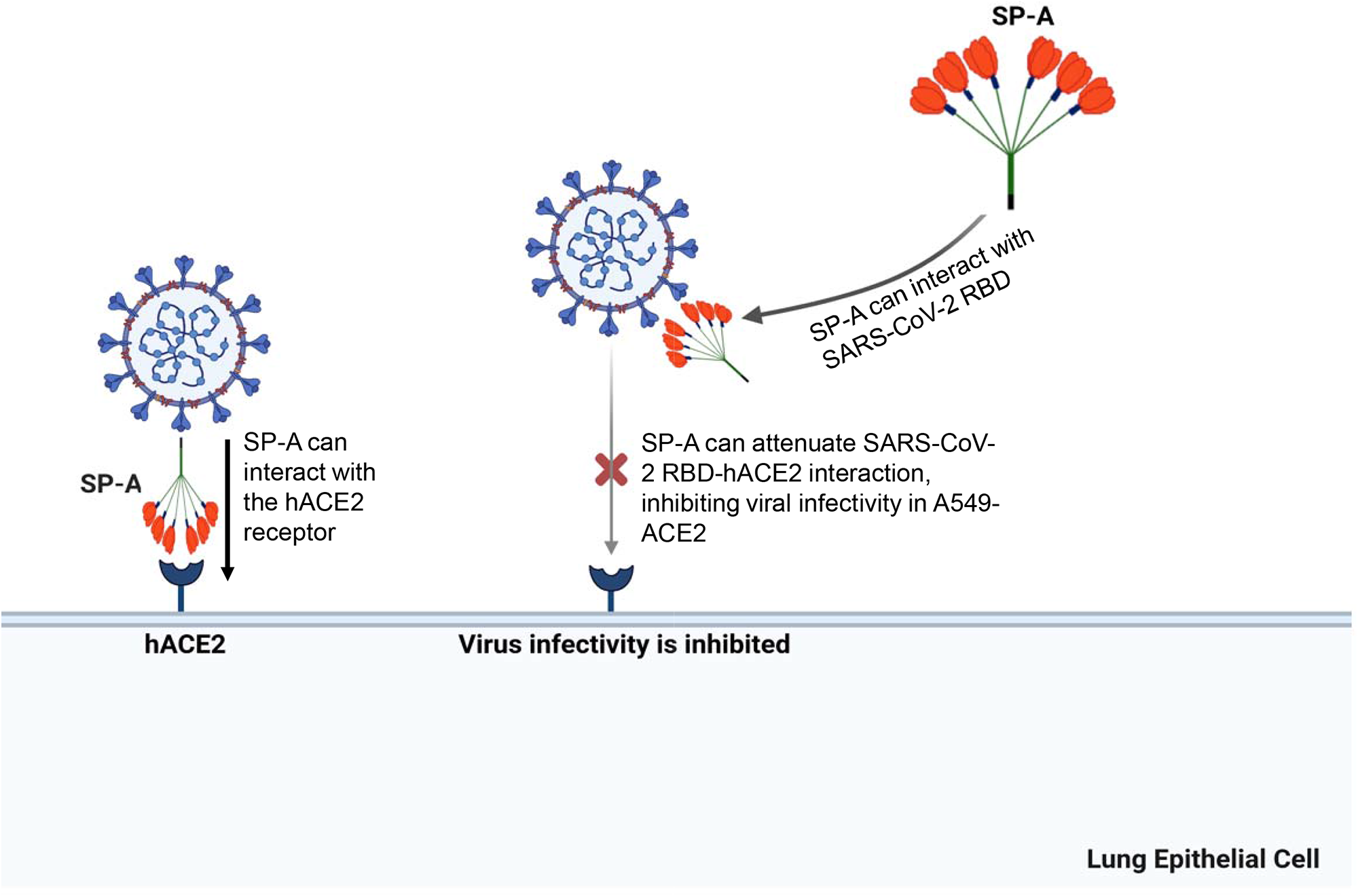
A diagram of SP-A interaction with S protein and human ACE2 receptor resulting in a significant decrease in SARS-CoV-2 infectivity. The SARS-CoV-2 spike protein can be recognized by human SP-A resulting in viral particle aggregations and reduced interactions of virus and host cell ACE2 receptors. Human SP-A also directly interact with human ACE2 receptor, which can inhibits the binding of SARS-CoV-2 RBD to hACE2, and subsequently diminishing viral entry and infectivity in susceptible host cells.

## ACKNOWLEDGMENTS

The authors thank all COVID-19 patients and healthy individuals who kindly provided saliva specimens used for this study. We would like to appreciate Dr. Joanna Floros for her generous support and encouragements to this project, Dr. Gary Chan for his experimental suggestions and Mr. Reuben Onwe for his critical reading and editing. A549-ACE2 (human lung adenocarcinoma epithelial cell overexpressing human ACE2) and Vero E6 cells were obtained through BEI Resources, NIAID, NIH.

## Notes

This work was supported by NIH R01HL136706, R21AI171574, the NSF research award (1722630), and the Richard and Jean Clark Pediatric Research Fund (to GW); and by the NIH 1R21AI14932101, 3R21AI149321-01S1, 1R01AI148446-01A1 (to HJ)

### Competing Interest Statement

The authors have declared no competing interest.

